# High-fidelity optical diffraction tomography of live organisms using non-toxic tunable refractive index media

**DOI:** 10.1101/2022.05.18.492426

**Authors:** Dohyeon Lee, Moosung Lee, Haechan Kwak, Youngseo Kim, Jaehyu Shim, Jik Han Jung, Wei-Sun Park, Ji-Ho Park, Sumin Lee, Yongkeun Park

**Affiliations:** Department of Physics, Korea Advanced Institute of Science and Technology (KAIST), Daejeon 34141, South Korea; KAIST Institute for Health Science and Technology, KAIST, Daejeon 34141, South Korea; Department of Bio and Brain Engineering, KAIST, Daejeon 34141, South Korea; Tomocube Inc., Daejeon 34109, South Korea

## Abstract

Optical diffraction tomography (ODT) enables the three-dimensional (3D) refractive index (RI) reconstruction. However, when the RI difference between a sample and a medium increases, effects of light scattering become significant, preventing the acquisition of high-quality and accurate RI reconstructions. Herein, we present a method for high-fidelity ODT by introducing non-toxic RI matching media. Optimally reducing the RI contrast enhances the fidelity and accuracy of 3D RI reconstruction, enabling visualization of the morphology and intra-organization of live biological samples without producing toxic effects. We validate our method using various biological organisms, including *C. albicans* and *C. elegans*.

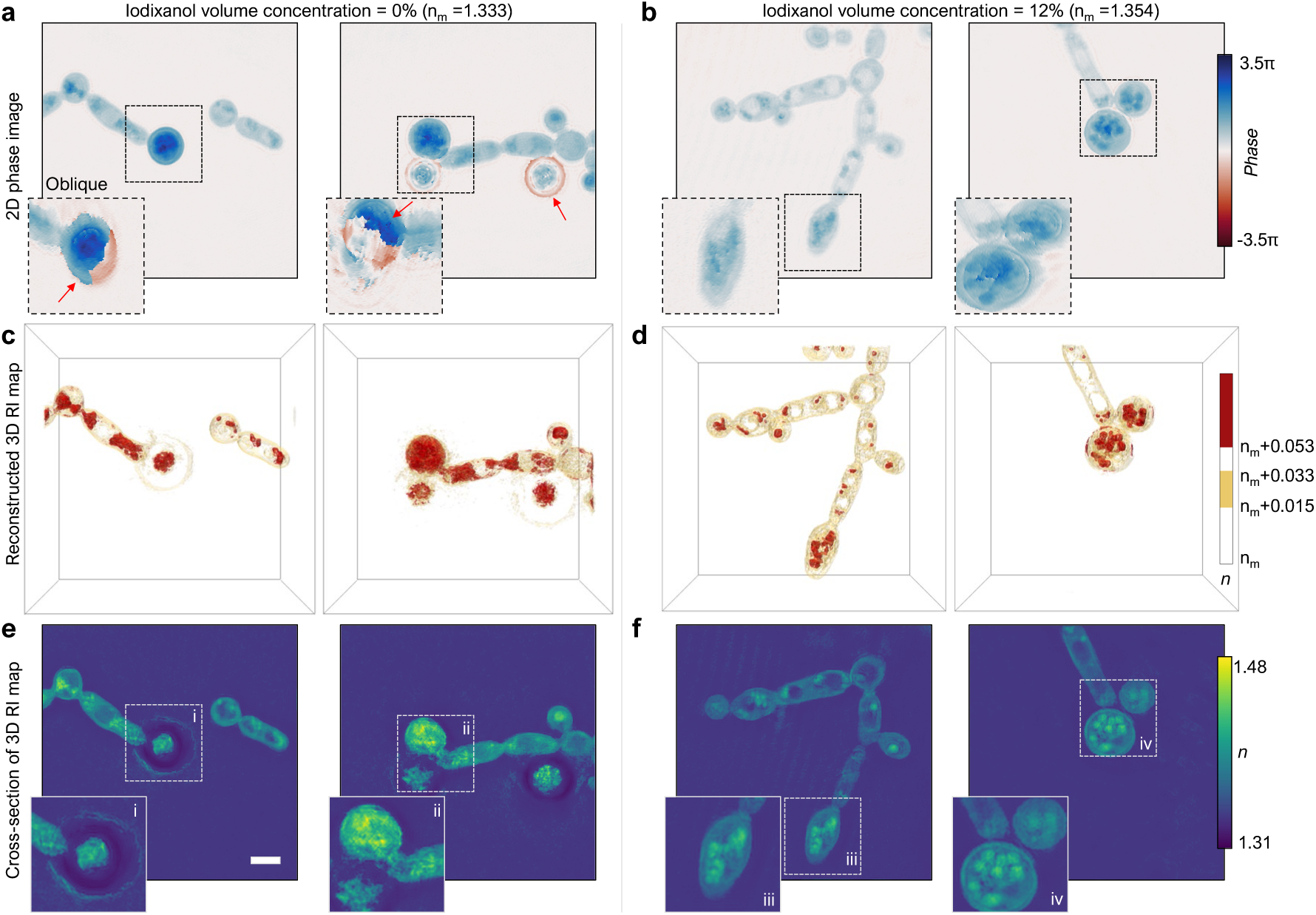

## 1. Introduction

Imaging of biological organisms such as nematodes is indispensable for understanding various biological processes, including genetic mutations, evolution, pathophysiology, and longevity [1]. Although various microscopic techniques have been utilized to study nematodes [2], the use of fluorescent proteins or staining dyes imposes several limitations, such as phototoxicity, photobleaching, and genetic mutations [3].

Recently, quantitative phase imaging (QPI) techniques have been employed for imaging and analyzing the dynamics of biological organisms owing to their label-free and quantitative imaging capabilities [4-9]. QPI uses the refractive index (RI) distribution of a sample as an intrinsic imaging contrast and enables three-dimensional (3D) imaging of live unlabeled biological samples [10-13]. However, unlike thin biological cells or microorganisms [11, 12, 14, 15], the large sizes and high RI values of nematodes result in technical challenges for QPI. Large optical phase delay variations in nematodes generate significant phase-wrapping issues, deteriorating the imaging quality of 2D QPI techniques. A high RI contrast between the sample and the surrounding medium results in multiple light scattering, which violates the assumption of weak scattering conditions. Thus, the 3D RI reconstruction of nematodes suffers from artifacts and poor image quality.

To mitigate the limited reconstruction fidelity resulting from large RI contrasts, the RI difference between a sample and medium can be reduced; this is analogous to impedance matching by ultrasound gels in medical ultrasound imaging [16, 17]. Commercially available refractive-index-matching media exist. However, most of these are prepared using mixtures of carcinogenic organic solvents that are incompatible with biological studies [18]. Alternatively, an aqueous solution of glycerol or sugar may be used for index matching [19, 20]. However, this solution alters various physiologically significant conditions such as osmolality, subsequently rendering it unviable for living cells.

In this study, we demonstrate enhancement of the fidelity of 3D QPI using nontoxic tunable RI media. By supplementing iodixanol as a non-toxic RI matching medium, the issue related to RI mismatch in imaging live organisms was alleviated, and tomographic RI distributions of organisms were clearly and precisely retrieved. Iodixanol displayed an osmolarity similar to that of typical vertebrate cell culture media, and the linearity between the RI and density was used to tune the RI of aqueous solutions between 1.333 and 1.429 [21]. Further, we validated the label-free tomographic imaging of live *C. albicans* and *C. elegans*. We envision that the proposed simple and cost-effective solution for imaging large organisms may broaden the applicability of QPI in the study of multicellular organisms.

## 2. Methods

### 2.1 Optical diffraction tomography

To acquire RI tomograms of live biological samples, we used an optical diffraction tomography (ODT) setup (HT-1H; Tomocube Inc., South Korea) [Fig. 1a]. ODT is a 3D QPI technique based on laser interferometric microscopy with a capability of varying illumination angles [22]. From the measured multiple 2D interferograms of a sample, its 3D RI tomogram can be reconstructed by applying the phase-retrieval algorithm and solving the inverse scattering equation [23].

**Fig. 1.**
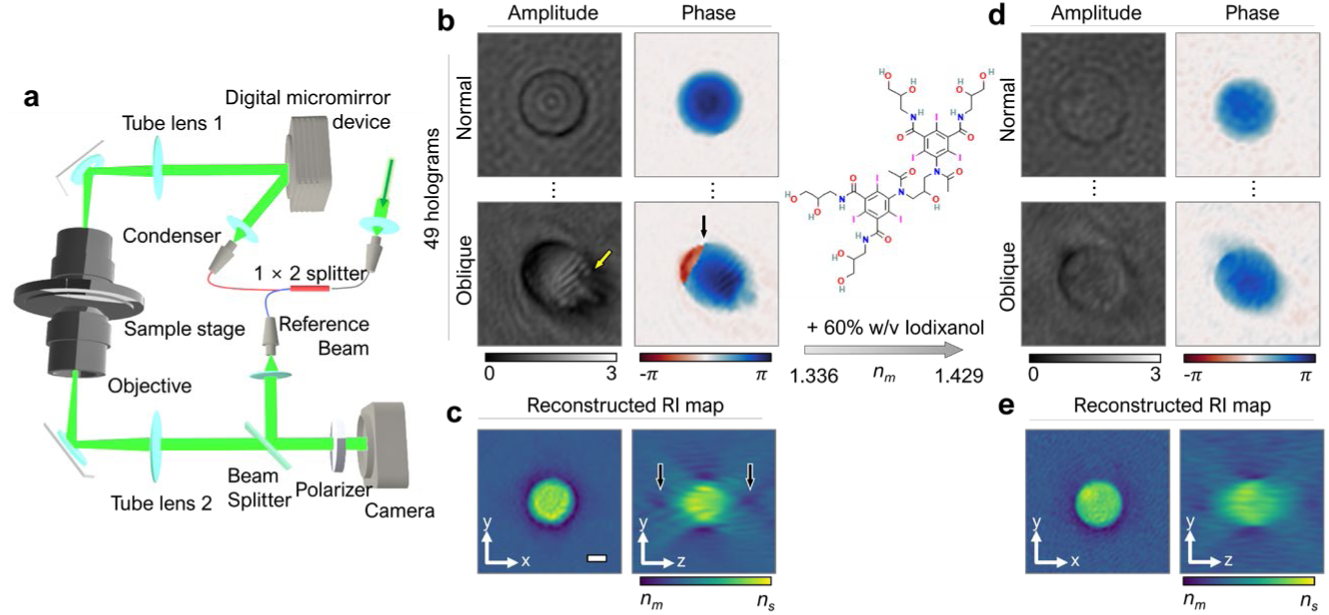
a, Schematics of an ODT system. **b–c**, Amplitude and phase images retrieved from holograms of a poly(methyl methacrylate) (PMMA) bead in deionized water, obtained at various illumination angles, from which a 3D RI map was reconstructed. **d–e**, Corresponding results of a PMMA bead in a 60% (w/v) iodixanol solution.

The setup was based on an off-axis Mach–Zehnder interferometric microscope equipped with a digital micromirror device (DMD) [24]. A beam from a coherent light source (*λ* = 532 nm in vacuum) was split into sample and reference arms using a 1 × 2 fiber splitter. A DMD (DLP6500FLQ, Texas Instruments, USA) systematically controls the illumination angle of the beam impinging onto a sample [25, 26]. To magnify and focus the light, the sample beam was passed through a 4-*f* telescopic imaging system consisting of a tube lens (tube lens 1, *f* = 250 mm) and a condenser lens (PLASAPO 60XW, Olympus Inc., Japan) with a numerical aperture (NA) of 1.2. The light diffracted from the sample was collected using an objective lens (UPLASAPO 60XW, Olympus Inc., Japan) and tube lens (Tube lens 2, *f* = 175 mm).

Both the sample and reference beams were combined using a beam splitter, and the corresponding spatially modulated interferogram was formed on an image plane, which was recorded using a camera (FL3-U3-13Y3M-C, FLIR Systems, Inc., USA). The polarizer was located in front of the camera to ensure a fringe contrast. For each tomogram measurement, 49 holograms were obtained at various illumination angles, which took 0.5 s.

### 2.2 3D RI reconstruction

From the measured multiple 2D holograms of a sample, a 3D RI tomogram was reconstructed by inversely solving the Helmholtz equation with the Rytov approximation, which assumes weak scattering [22]. From each hologram obtained with a specific illumination angle, an optical field image containing both the amplitude and phase information was obtained using a field retrieval algorithm [Fig. 1b]. Multiple 2D optical field images were mapped onto a 3D Fourier space according to the Fourier diffraction theorem [22, 27]. The theoretical spatial resolutions of the ODT system used were 110 and 360 nm in the lateral and axial directions, respectively [28].

Owing to the small NAs of the condenser and objective lenses, side-scattering signals cannot be recorded. This results in deterioration of the image quality of the reconstructed tomogram, including poor axial resolution and underestimation of the RI values. To compensate for this issue, known as the missing cone problem, we used total variance regularization accelerated by the fast iterative shrinkage and thresholding algorithm (TV-FISTA) with a non-negativity constraint [29, 30]. Detailed information on the field retrieval [31], reconstruction algorithm [23], and regularization [30] can be found elsewhere.

### 2.3 Preparation of live specimens

*Candida albicans* (*C. albicans*) strain CEC2018 was cultured at 25°C for 48 h in a liquid potato-carrot-bile medium to observe chlamydospore formation. Twenty microliters of the culture was homogeneously mixed with 5 µL of a 60% iodixanol solution by gentle pipetting. To observe *Caenorhabditis elegans* (*C. elegans*) in the presence of iodixanol, worms on a nematode growth medium (NGM) agar plate were transferred into a liquid Luria-Bertani (LB) medium. The concentration of iodixanol was adjusted by mixing a correlated volume of Optiprep and the worm suspension or by serial dilution of the worm mixture using the liquid LB medium. The mixture was immediately mounted onto a TomoDish (Tomocube Inc., Republic of Korea) by using a #1.5H coverslip (Paul Marienfeld GmbH & Co. KG, Germany) to acquire the RI tomograms.

### 2.4 Preparation of plastic microspheres

A drop of poly(methyl methacrylate) (PMMA) beads (Cat No. 86935, Sigma-Aldrich, USA) with diameters of 3 μm was placed onto a #1.5H coverslip (Marienfeld, Germany) without dilution, after which it was dried. After complete drying, a drop of the surrounding medium was gently dropped onto the sample and then covered with a #1.5H coverslip before measurements.

## 3. Results

### 3.1 Improvement of the ODT image quality by RI matching using iodixanol

To demonstrate the effects of RI matching using iodixanol on the ODT image quality, we measured PMMA microspheres with diameters of 3 μm [Fig. 1b–1e]. When the PMMA in deionized (DI) water is measured, the amplitude and phase images are well retrieved under normal illumination [Fig. 1b]. However, under oblique illumination, the amplitude map demonstrates diffraction patterns (yellow arrow in Fig. 1b), and the phase map exhibits a wrapped phase (black arrow in Fig. 1b). The reconstructed RI tomogram presents artifacts, including the halo effect adjacent to the sample boundary and the diffraction patterns inside the bead and along the axial direction (black arrows in Fig. 1c).

The PMMA bead sample used was on the borderline between the weak scattering and multiple scattering regimes. The RI difference between the bead and the surrounding medium was Δ*n = n*_*s*_ *– n*_*m*_ *=* 1.493 *–* 1.333 = 0.160, and the corresponding optical phase delay for the normal illumination was Δ*ϕ* = 2π/*λ* · Δ*n* · *h* ≈ 1.8π, which is lower than 2π. For oblique illumination, the optical phase delay can be greater than 2π, resulting in phase wrapping, and the effects of multiple scattering inside the sample are not negligible. This situation becomes more significant when the size of the sample becomes larger, or the RI difference between the sample and the surrounding medium increases. Thus, an important assumption for the ODT algorithm – the weak scattering condition – is violated, and precisely retrieving the RI tomogram becomes challenging.

Following this, we repeated the measurement of the PMMA bead in the 60% iodixanol (w/v) medium. In this case (Fig. 1d), the amplitude and phase images of the bead are retrieved under both normal and oblique illumination. No strong diffraction patterns or failures during phase unwarping are observed. This result is consistent with recent reports, wherein the addition of iodixanol improved the image quality of 2D QPI [32-35]. The reconstructed 3D RI tomogram of the PMMA beads in the presence of iodixanol demonstrated that the artifacts shown in Fig. 1c were alleviated, and the diffraction patterns inside the bead and along the axial direction were significantly reduced.

### 3.2 Quantification of the RI values of the Iodixanol medium

To verify the RI tunability of the iodixanol medium, the RI values of the medium at various concentrations were quantified. A 60% (w/v) iodixanol solution (Optiprep; Cat No. D1556, Sigma-Aldrich, USA) was diluted with distilled water to prepare 0%, 10%, 20%, 30%, 40%, 50%, and 60% iodixanol media. The RI of each medium was measured using a refractometer (Atago™ R-5000; ATAGO Co., Ltd., Tokyo, Japan) [Fig. 2]. The results demonstrate a linear relationship between the measured RI and iodixanol concentration over the range of 0% to 60%.

**Fig. 2.**
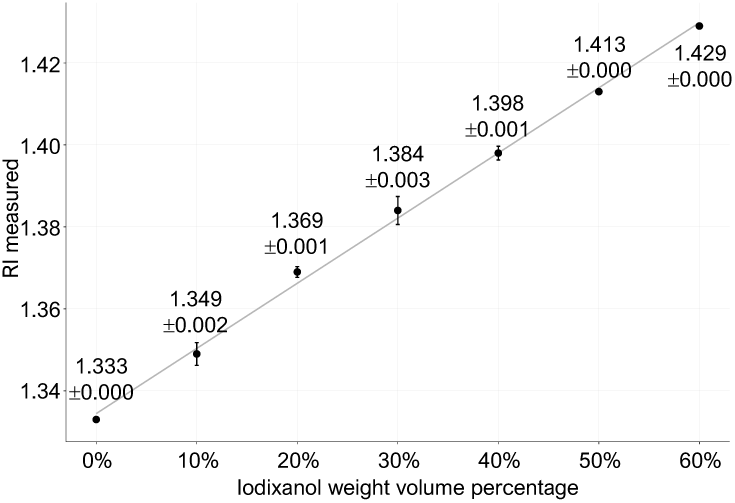
RI values of iodixanol solutions with various concentrations.

### 3.3 Quantification of the reconstruction quality using a reference PMMA bead sample

To demonstrate the improvement in ODT image quality by RI matching, PMMA microspheres with diameters of 3 μm were measured with various concentrations of iodixanol (0%–60%). After mixing the PMMA beads and iodixanol medium, the individual microspheres were measured using the ODT setup (Methods).

The measured RI tomograms of PMMA microspheres in the media with various RI values are presented in Fig. 3. An increase in the iodixanol concentration leads to proper evaluation of the RI around the boundary of the beads. Without any index-matching process (0% iodixanol), the reconstructed RI tomograms exhibit artifacts, including halo and squeezing artifacts [Fig. 3a]. The former can be attributed to the limited axial resolution caused by the missing cone problem, and the latter can be attributed to the failure in phase unwrapping, which is important for high-fidelity RI reconstruction. The addition of iodixanol mitigated squeezing artifacts by reducing the severe RI contrast, which impeded accurate RI reconstruction. Artifacts can also be observed in the *y-z* cross-sections. When the RI was not matched, halo and squeezing artifacts were dominant, whereas when RI was matched (60%), the image quality was significantly improved. In addition, diffraction patterns were suppressed when the RI contrast was matched.

**Fig. 3.**
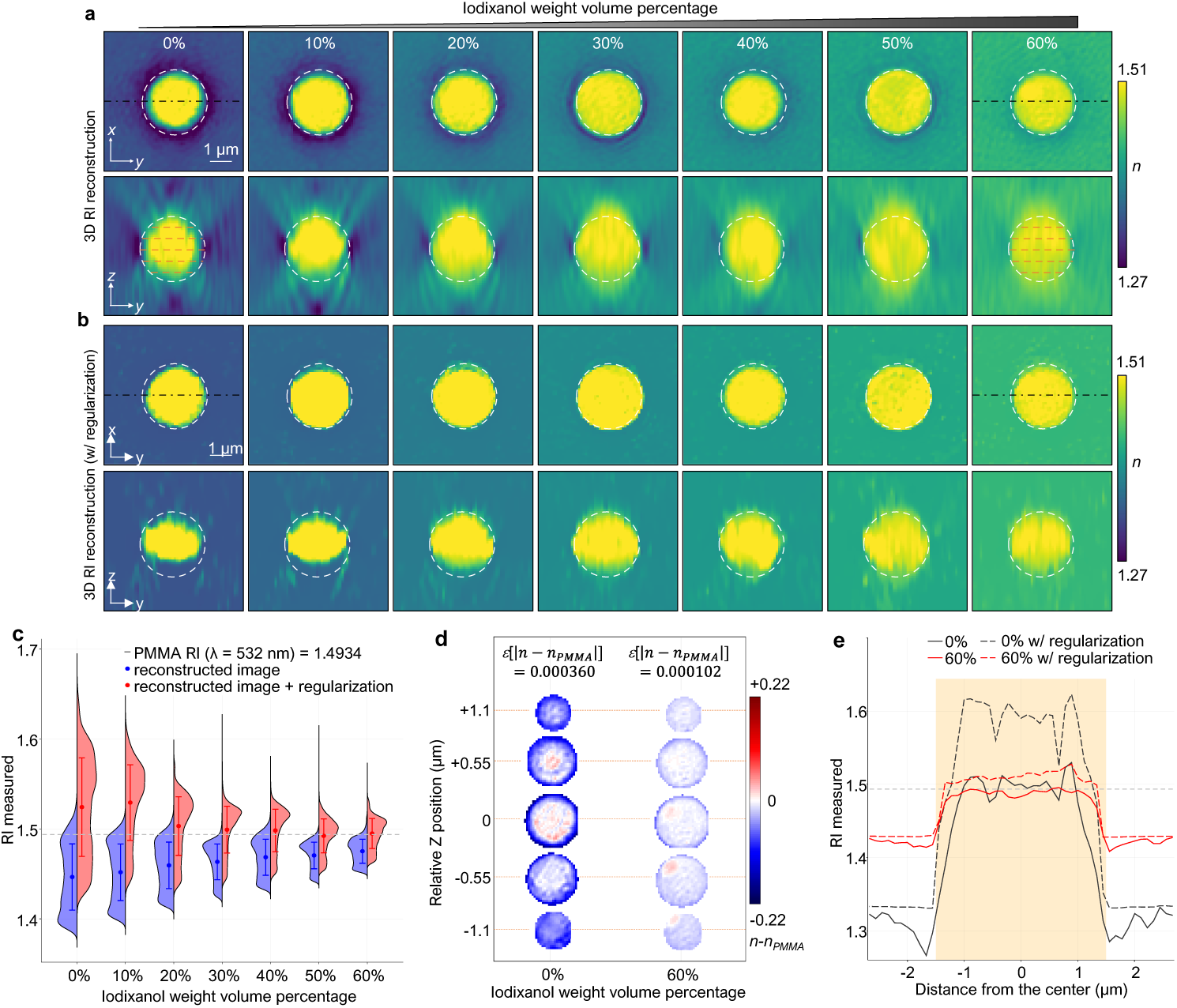
**a**, Cross-sectional images of a reconstructed 3D RI tomogram of PPMA beads in iodixanol solutions with various concentrations. **b**, Results after applying the regularization algorithm. **c**, Statistics of the retrieved RI values in **a** and **b. d**, Mean absolute error maps between the reconstructed RI and expected values at various axial positions. **e**, Profiles of RI values along the *y*-directional lines at the focal plane (the black dash-dotted lines in **a** and **b**).

We then tested the effects of the regularization algorithm on the measured RI tomograms under various RI mismatch conditions. The regularization algorithm is used to compensate for the missing cone problem, which results from the small NAs of the condenser and objective lenses. We applied the TV-FISTA regularization algorithm to the reconstructed 3D RI tomograms of the PMMA beads [Fig. 3a], and the results are presented in Fig. 3b. It is well known that the application of a regularization algorithm mitigates the issue of axial elongated tomographic reconstruction. However, when the RI contrast is high, that is, at low iodixanol concentrations, the reconstructed tomograms of the microspheres exhibit axially compressed ellipsoidal shapes along the axial direction. In contrast, when the RI is matched, the reconstructed tomogram demonstrates spherical shapes in all directions.

To quantitatively analyze the values of the reconstructed RI tomograms, we compared the reconstructed RI distributions of the PMMA microspheres with the expected values (*n* = 1.4934) [Fig. 3c]. As the RI contrast between the sample and medium is matched using iodixanol, the accuracy and precision are enhanced. Before applying the regularization algorithm (blue histograms in Fig. 3c), the RI values were underestimated owing to the missing cone problem. However, after applying the regularization algorithm, the mean value of the reconstructed RI tomograms decreased as the concentration of iodixanol increased.

To investigate the distribution of errors, error maps, calculated as the difference between the measured and expected RI values, *n* – *n*_*PMMA*_, are presented at various axial positions [Fig. 3d]. At an iodixanol concentration of 60%, the absolute mean value of error, *ε*(|*n* – *n*_*PMMA*_|), decreases by a factor of 3.529. In addition, correction of the RI values is dominantly observed at the boundaries of a sample, suggesting that the halo effects resulting from multiple scattering are significantly decreased by matching the RI contrast. Consequently, the RI profile of a PMMA microsphere matches well with the expected distributions when a 60% iodixanol medium is used with the regularization algorithm [Fig. 03e].

### 3.4 Quantification of the reconstruction quality using a reference PMMA bead sample

To verify the applicability of the proposed method, we measured *C. albicans*, a pathogenic yeast that causes life-threatening diseases in humans through the skin or mucosal membranes. Imaging of the death process of *C. albicans* upon antifungal treatment is important for the development of new antifungal agents. Several studies have monitored antifungal activity by observing the morphological alteration of *C. albicans* cells in 3D cultures [36, 37]. However, a quantitative analysis of internal phenotypic alterations, such as protein density, has not been completely performed. Although soft X-ray tomography can provide high-resolution images of subcellular changes upon antifungal treatment [38], observations of the dynamic process of live yeast are complicated in cryogenic X-ray imaging. Recently, ODT has been used to study the dynamics of *Yarrowia lipolytic* through internal epidemiological dynamics [39]. One of the challenges encountered in imaging *C. albicans* is the high RI value of yeasts, which makes it difficult to retrieve 2D phase and 3D RI tomogram images.

Using the proposed method, we measured both the 2D phase and 3D RI tomograms of *C. albicans* [Fig. 4]. In the conventional approach without iodixanol, 2D phase images exhibit strong artifacts owing to the failure of the phase unwrapping algorithm [Fig. 4a]. These artifacts are dominant in chlamydospores – spherical and thick-walled spores with dense high-RI internal structures. The proposed method resolves these issues and demonstrates a clear 2D phase [Fig. 4b]. For a 12% (w/v) iodixanol solution, phase maps display a smooth spatial change in phase images under both normal and oblique illuminations.

**Fig. 4.**
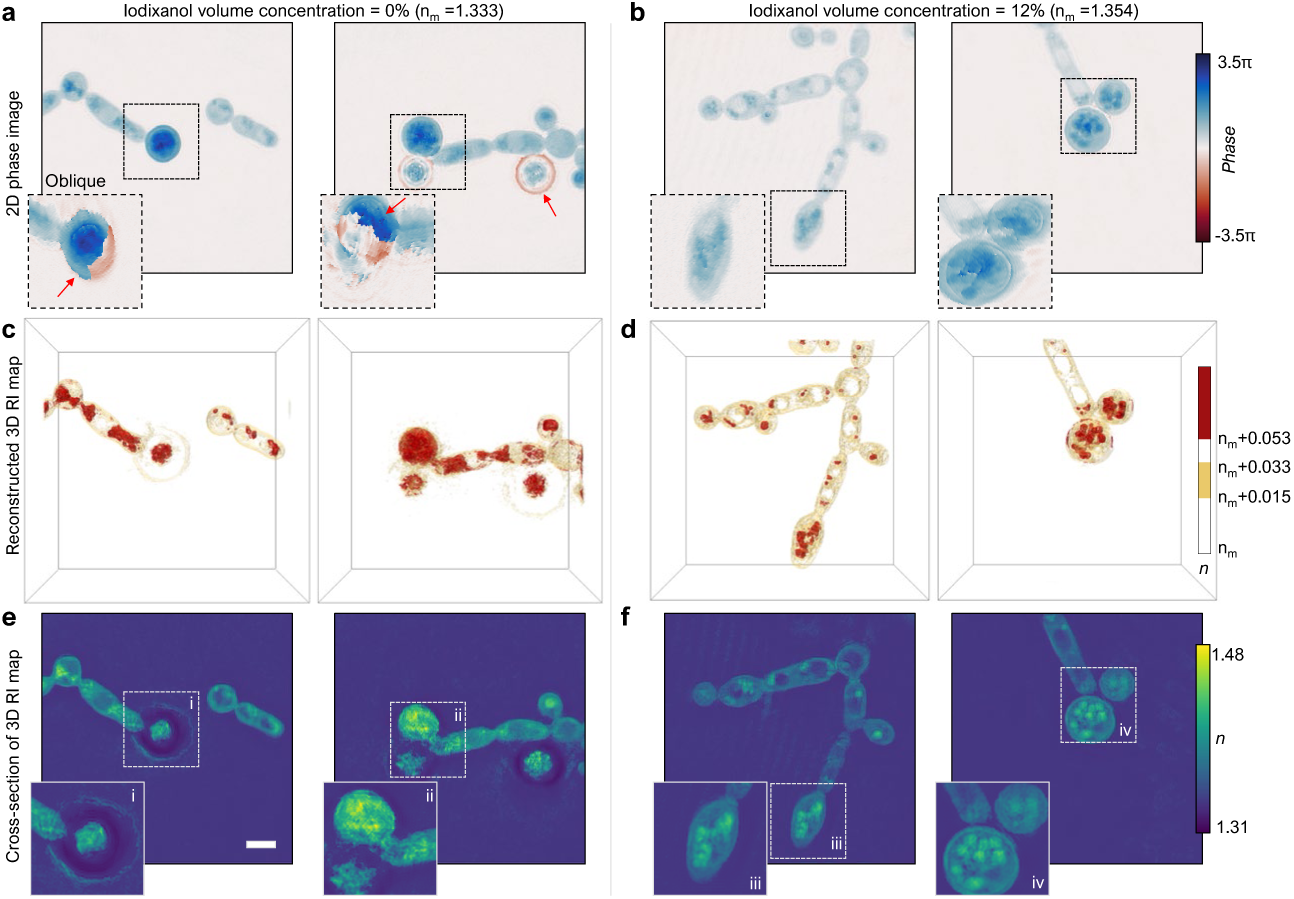
**a–b**, 2D phase images of *C. albicans* in DI water (**a**) and in a 12% (w/v) iodixanol solution (**b**). **c–d**, Reconstructed 3D RI tomogram of *C. albicans* in (**a**) and (**b**). **e–f**, Cross-sectional images of the RI tomogram in the *x-y* planes in (**c**) and (**d**).

Matching of the RIs between *C. albicans* and the surrounding media using iodixanol enhances the image quality of the 3D RI tomograms [Fig. 4c–f]. Owing to the high RI contrast between the sample and surrounding media, the reconstruction algorithm generates several artifacts. For example, a chlamydospore exhibits a RI smaller than that of the surrounding medium, which is not physically possible (inset i). In addition, the reconstructed tomogram of another chlamydospore presents blurred internal structures (inset ii). In contrast, the proposed method generates high-fidelity 3D structures of *C. albicans*. The reconstructed RI tomograms clearly present the walls and internal structures of the pseudohyphae and chlamydospores with high resolution. In particular, the dense structures of chlamydospores are reconstructed without artifacts (insets iii and iv).

### 3.3 3D RI tomograms of C. Elegans

To further demonstrate the applicability of the proposed method for studying nematodes, the pharynx region of *C. elegans* was measured [Fig. 5]. Previously, label-free imaging of *C. elegans* has been investigated owing to its importance in the study of cellular differentiation and developmental processes in intact organisms. However, the nature of multicellular eukaryotic organisms and their high internal RI distributions cause light scattering [4, 40].

**Fig. 4.**
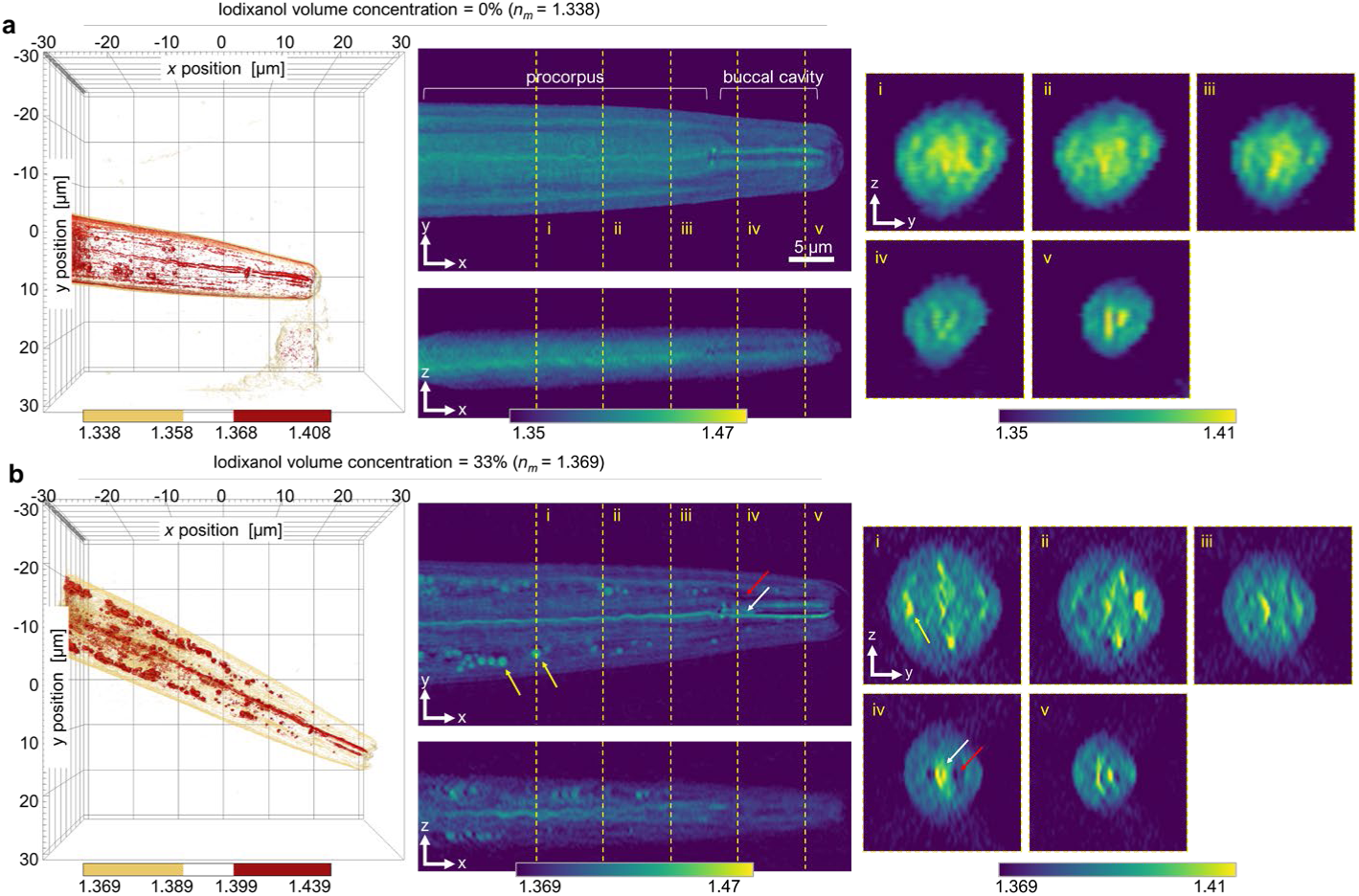
**a–b**, Reconstructed 3D RI tomogram of the digestive system of *C. elegans* measured in DI water (**a**) and in a 20% (w/v) iodixanol solution (**b**). The imaged area includes the procorpus and buccal cavity region. Each RI slice was captured alongside the esophagus. The subsets (i)–(v) present cross-sectional views along the *y-z* plane corresponding to the yellow dashed lines in **(a)** and **(b)**.

When a *C. elegans* worm is measured in a liquid LB medium, the overall shape of the worm and internal structures, such as the procorpus or buccal cavity, can be observed [Fig. 5a]. However, owing to the high RI contrast between the sample and the surrounding medium, details pertaining to small internal structures are difficult to identify at a high resolution. When the RI of the surrounding medium is matched by adjusting the concentration of iodixanol to 20%, multiple granular structures, as well as the outlines of the procorpus or buccal cavity, can be observed [Fig. 5b]. The sizes, shapes, and high RI values (*n* > 1.4) of the granular structures (yellow arrows) suggest that they are lipid droplets [41, 42], which are known to be located in the outer region of the procurpus [43]. This suggests that the structure indicated by the red arrow is an endometrial flap, and the boundary between the stoma and procorpus is indicated by the white arrow.

## 4. Conclusion

Herein, we report that the ODT image quality of optically thick samples can be improved by adding iodixanol, a non-toxic tunable RI medium. This enhancement in image fidelity is enabled by RI matching between a biological sample and the surrounding media, resulting in a decrease in scattering effects. In the current study, we validated this method by imaging various nonbiological and biological organisms, including PMMA microspheres, *C. albicans*, and *C. elegans*. This method is robust, facile, biocompatible, and potentially adaptable to a wide variety of studies, wherein 3D QPI is used to image thick biological samples [44-49], such as spheroids, organoids, embryos, and tissues. Furthermore, the proposed method can be used for quantitative imaging of nonbiological samples with high RI contrasts, including microplastics and bacteria [14, 50].

Although the proposed method significantly improves the image quality of ODT, multiple scattering effects may not be completely suppressed owing to various unresolved issues. For example, issues related to strongly inhomogeneous distributions of RI inside biological samples or distinctly high RI regions inside a sample may be completely addressed by index matching between the sample and the surrounding medium. The unaddressed multiple scattering issues can be resolved using a reconstruction algorithm that considers multiple light scattering [51-53]. In addition, machine-learning approaches can be utilized to further suppress multiple light-scattering effects [54].

The proposed method is straightforward, facile, and cost-effective and will enable label-free quantitative analysis of the morphology and dynamics of complex biological systems. The label-free imaging capability of ODT can demonstrate synergetic effects in combination with the use of non-toxic tunable RI media for imaging large biological samples over a long time. Iodixanol has been proven to be biocompatible in cell imaging [21], and the label-free nature of ODT enables long-term monitoring of cells that are implanted into humans [55-57]. Given the simple and biocompatible nature of the proposed method, it could potentially be used for the study of developmental biology or regenerative medicine [58, 59]. We envision that this synergetic approach can have far-reaching applications, possibly in conjunction with emerging single-cell profiling in stem cell or immune therapy.

## Acknowledgments

*Candida albicans. C. elegans* and *Candida albicans* were kindly provided from Jun Kim, Ph.D. (Seoul National University) and Sophie Bachellier-Bassi (Institute Pasteur), respectively.

## Data availability

The source code and data can be provided upon reasonable requests.

